# Human ovarian ageing is characterized by oxidative damage and mitochondrial dysfunction

**DOI:** 10.1101/2023.01.31.525662

**Authors:** Myrthe A.J. Smits, Bauke V. Schomakers, Michel van Weeghel, Eric J.M. Wever, Rob C.I. Wüst, Frederike Dijk, Georges E. Janssens, Mariëtte Goddijn, Sebastiaan Mastenbroek, Riekelt H. Houtkooper, Geert Hamer

## Abstract

Human ovarian ageing encompasses the age-related decline in female fertility. Oxidative stress and mitochondrial dysfunction in oocytes are suggested as causal, but corroborating evidence is limited. Using immunofluorescence imaging on human ovarian tissue, we found oxidative damage by protein and lipid (per)oxidation at the primordial follicle stage. Additionally, using comprehensive metabolomics and lipidomics, a cohort of 150 human germinal vesicles and metaphase I oocytes and 15 corresponding cumulus cell samples displayed a shift in glutathione to oxiglutathione ratio and depletion of phospholipids. Age-related changes in polar metabolites suggested a decrease in mitochondrial function, as demonstrated by NAD^+^, purine and pyrimidine depletion, while glycolysis substrates and glutamine accumulated with age. Oocytes of advanced maternal age likely used alternative energy sources like glycolysis and the adenosine salvage pathway, and possibly increased ATP production in cumulus cells. These findings indicate that oocytes of advanced maternal age suffer from oxidative damage and mitochondrial dysfunction.

**Graphical abstract:** 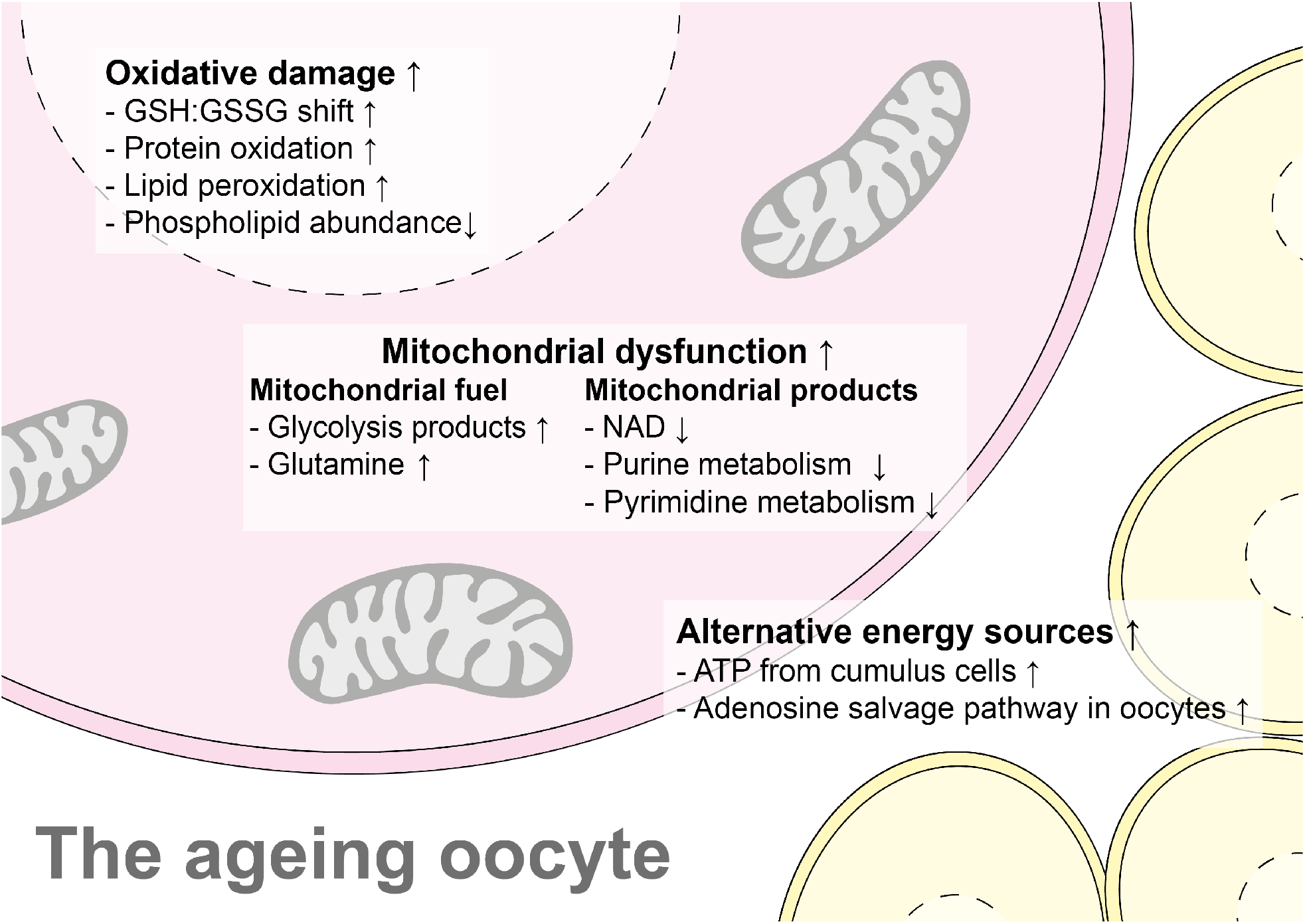

## Introduction

Chances of reproductive success in humans decline with increasing female age, a phenomenon known as ovarian ageing. Women of advanced maternal age (A>35 years old) have lower chances of natural conception (Eijkemans et al., 2014), as well as lower success rates of assisted reproductive techniques and increased chances of pregnancy loss (CDC, 2016, Schieve et al., 2003). Embryo transfers have led to pregnancy in 9.8% of women over 40 years old using their own oocytes, but when women of this age group were given younger donor oocytes, their pregnancy success rate was 48.7% (CDC, 2016). The age-related decline in reproductive success appears therefore largely dependent on the age-related decrease in oocyte quality. The driving force behind the decrease in oocyte quality with increasing female age is not yet fully understood and different hallmarks of somatic ageing have been studied in ovarian ageing (Chiang et al., 2020, Polonio et al., 2020, Kirkwood, 1998). One of the suggested mechanisms responsible for ovarian ageing is the accumulation of reactive oxygen species (ROS)-induced damage, which has been associated with mitochondrial dysfunction (May-Panloup et al., 2016, Lefkimmiatis et al., 2020, Wang et al., 2013, Hekimi et al., 2011). Mitochondrial dysfunction can in turn result in failure of multiple cell organelles, apoptosis and cellular senescence (Wang et al., 2013, Vasileiou et al., 2019).

Female meiosis initiates prenatally in the fetal gonads but arrests at the end of the first meiotic prophase just before the first meiotic division would otherwise occur (Faddy et al., 1992). Already before birth, these so-called dictyate arrested oocytes reside in primordial follicles, consisting of dictyate arrested oocyte and supporting granulosa cells, which are then waiting in the ovary to be stimulated for maturation, a period that can span 40+ years (Faddy et al., 1992). During this period of cell cycle arrest the primordial follicles maintain basic metabolism to stay alive, while being exposed to environmental factors. Even in a situation where only basic metabolism is needed to maintain cell homeostasis for a long time, cells can get exposed to ROS-induced oxidative damage and mitochondrial dysfunction (Ahmed et al., 2019, Cinco et al., 2016).

During the reproductive years of life in women, under the influence of follicle stimulating hormone, follicles start to mature in the ovary, a process called folliculogenesis. Folliculogenesis starts at the primordial follicle stage. The primordial follicle consists of a germinal vesicle (GV) oocyte with few surrounding flattened granulosa cells. This is the stage at which follicles reside from birth onwards until they are stimulated for maturation (Figure 1A). During the primary follicle stage, the surrounding granulosa cells increase in number and become more cubically shaped (Figure 1B). In the secondary follicle stage, the oocyte increases in size and surrounding granulosa cells form multiple layers (Figure 1C). During the antral follicle stage, the final stage of folliculogenesis before ovulation, a fluid filled cavity develops between different layers of granulosa cells (Figure 1D). After ovulation, the oocyte resumes meiosis I, through the metaphase I (MI) stage until it finally reaches the metaphase II stage where it arrests again until fertilization. The granulosa cells still associated with these oocytes are being referred to as cumulus cells.

**Figure 1.**
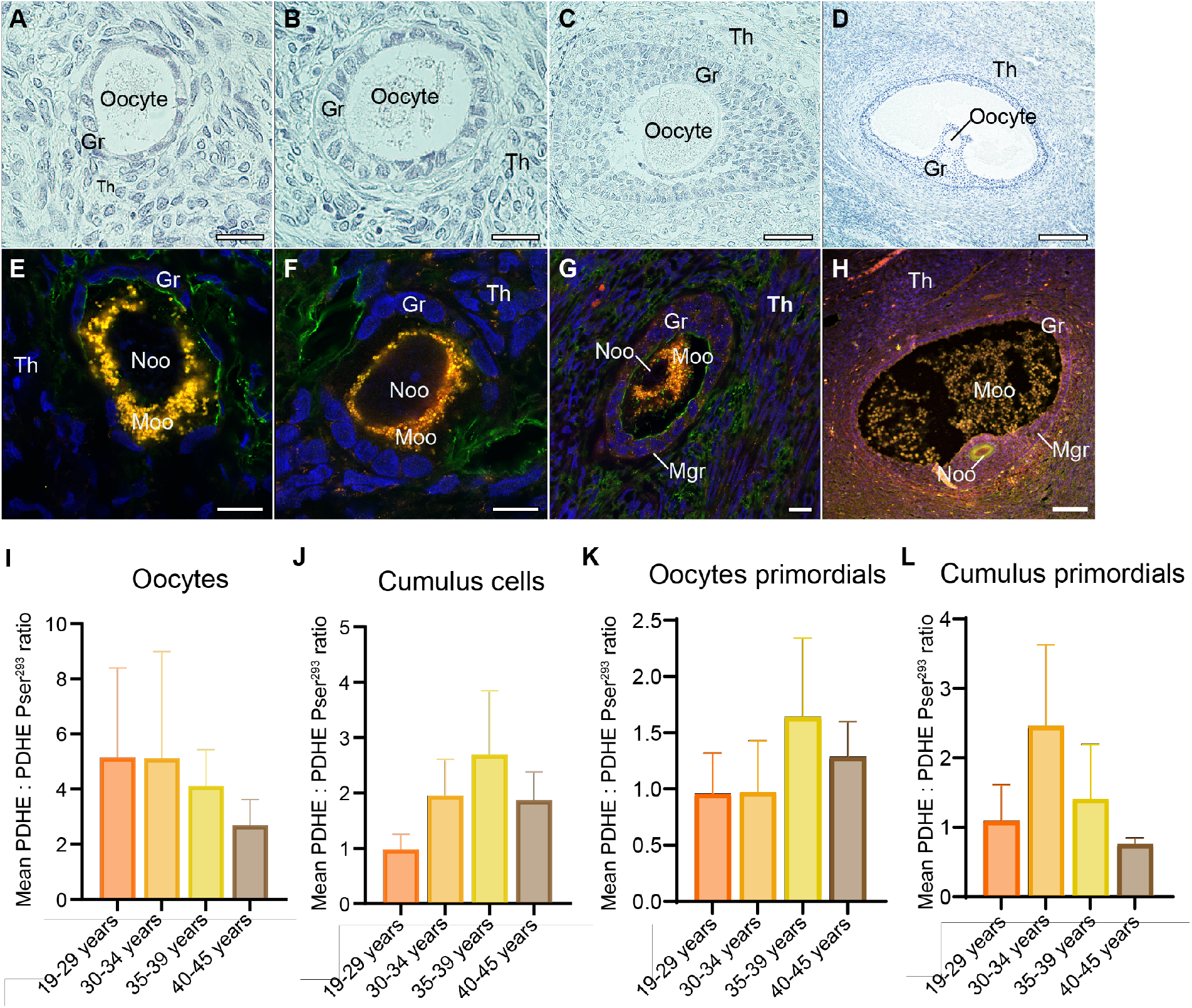
Pyruvate dehydrogenase (PDH) activation status in different stages of folliculogenesis of human oocytes. Upper row: staining of different stages of folliculogenesis. Lower row: Immunofluorescence staining of different stages of folliculogenesis. (A) Primordial follicle hematoxylin and eosin staining (scale bar 20μm). (B) Primary follicle hematoxylin and eosin staining (scale bar 20μm). (C) Secondary follicle hematoxylin and eosin staining (scale bar 50μm). (D) Antral follicle hematoxylin and eosin staining (scale bar 200μm). (E) Primordial follicle immunofluorescence staining (scale bar 10μm). (F) Primary follicle immunofluorescence staining (scale bar 10μm). (G) Secondary follicle immunofluorescence staining (scale bar 10μm). (H) Antral follicle immunofluorescence staining (scale bar 100μm). (I) Mean ratio of PDHE: PDHE1α pSer^293^ immunofluorescence staining as an indicator of oxidative phosphorylation activity per age group in oocytes. (J) Mean ratio of PDHE: PDHE1α pSer^293^ immunofluorescence staining as an indicator of oxidative phosphorylation activity per age group in cumulus cells. (K) Mean ratio of PDHE: PDHE1α pSer^293^ immunofluorescence staining as an indicator of oxidative phosphorylation activity per age group in oocytes from primordial follicles. (L) Mean ratio of PDHE: PDHE1α pSer^293^ immunofluorescence staining as an indicator of oxidative phosphorylation activity per age group in cumulus (granulosa) cells from primordial follicles. Gr = granulosa/cumulus cells, Th = theca cells. Channels: Red = PDHE, Yellow = phosphorylated PDHE (inactive PDH), Blue = DAPI (DNA), Green = Wheat germ agglutinin (WGA; membrane). Noo = nucleus oocyte, Moo = mitochondria oocyte, Mgr = mitochondria granulosa cells.

As a potential mechanism to limit ROS-induced damage in primordial follicles, it has been suggested that primordial follicles avoid using oxidative phosphorylation as an energy source and preferably rely on anaerobic glycolysis. However, recent evidence shows that oocytes already display signs of oxidative damage during the earliest stages of folliculogenesis, and mitochondria are already metabolically active (Van Blerkom, 2011, Jansen and de Boer, 1998, Bradley and Swann, 2019, Wang et al., 2020). During later stages of folliculogenesis, and even early stages of embryogenesis mitochondria become the primary energy source, and play an important role spindle formation (Hashimoto et al., 2017, Wilding et al., 2003). Signs of ROS-induced damage and mitochondrial dysfunction have been observed during these later stages of folliculogenesis and embryogenesis, signs of ROS-induced damage and mitochondrial dysfunction have been observed (Wang et al., 2017, May-Panloup et al., 2016, Pasquariello et al., 2019, Wang et al., 2021, Wilding et al., 2001). We postulate that ROS-induced damage and mitochondrial dysfunction may already occur in primordial follicles and thus play a major role in ovarian ageing.

In this study we investigated the activation status of pyruvate dehydrogenase, a key enzyme in oxidative phosphorylation, throughout all stages of folliculogenesis in paraffin embedded ovarian biopsies obtained from women of various ages. We subsequently investigated ROS-induced damage at the primordial follicle stage in these biopsies. Finally, we analyzed 90 human GV and 60 human (MI) oocytes and surrounding cumulus cells, collected during intracytoplasmic sperm injection (ICSI) treatment, using metabolomics and lipidomics analyses. We studied the role of cellular metabolism, and mitochondrial function in particular, in oocytes of different female ages. Using these different approaches, we found that older oocytes display all the investigated hallmarks associated with oxidative stress and mitochondrial dysfunction.

## Results

### All follicular stages, including the primordial follicles, display mitochondrial activity

To investigate possible age-related changes in human follicles, we collected formalin-fixed paraffin embedded ovarian tissue samples of 39 women of different ages who underwent surgery for non-malignant purposes. In hematoxylin eosin-stained tissue-sections, the presence of primordial, primary, secondary, and antral follicles was assessed (Figure 1A-D). If at least one follicular stage was present in a tissue section, the tissue was used for further super resolution immunofluorescence microscopy.

Since oxidative stress and mitochondrial dysfunction have often been implicated in matured oocytes of advanced maternal age (May-Panloup et al., 2016, Kasapoğlu and Seli, 2020), we decided to study the activation status of the mitochondrial gatekeeper enzyme pyruvate dehydrogenase (PDH) during all different stages of folliculogenesis. PDH is a major regulator of glucose metabolism, directing pyruvate to oxidative phosphorylation, rather than to anaerobic glycolysis. To this end, we used antibodies against the PDH subunit pyruvate dehydrogenase E1-alpha (PDHE1α), representing the total amount of PDH, in combination with antibodies against phosphorylated PDHE1α (PDHE1α pSer^293^), representing its inactive state. The active proportion of PDH was calculated by measurement and quantification of the PDHE1α to PDHE1α pSer^293^ fluorescent signal intensity ratios. The PDHE1α pSer^293^ staining, represented in yellow in Figure 1E-H, co-localized with the PDHE1α antibody, which is represented in red. To define the border between the oocyte and surrounding granulosa cells, we used the membrane marker Wheat Germ Agglutinin (WGA, represented in green), and 4’,6-diamidino-2-fenylindool (DAPI, represented in blue) to stain DNA (Figure 1E-H). Since the oocyte nucleus is very large in size, the DAPI staining more strongly visualizes the surrounding granulosa and theca cells. The PDH activation status, i.e. PDHE1α to PDHE1α pSer^293^ ratio, was calculated for all stages of folliculogenesis. We next compared how the PDH activation status changed in oocytes and granulosa cells with increasing female age. We included the following number of follicles into the analysis of the different age groups: 19-29 years, 14 follicles; 30-34 years, 9 follicles; 35-39 years, 14 follicles, and, 40-45 years, 22 follicles; at several follicular stages (supplementary data 1A. Although there was a trend towards a lower ratio and hence lower PDH activation status, when all follicles were analysed, no significant age-related change in PDH activation status was found in oocytes (p=0.612; Figure 1I) or granulosa cells (p=0.334) in regard to PDH activation status (Figure 1J). Because ovarian ageing occurs predominately at the primordial follicle stage, we additionally analysed oocytes (n =24) and granulosa cells (n =24) of this stage. At this follicular stage also, no significant age-related changes were observed in oocytes (p= 0.876, Figure 1K) or granulosa cells granulosa cells (p=0.859, Figure 1L). Negative controls showed no staining for PDHE1α and PDHE1α pSer^293^ (Figure S1A). Since no difference in PDH activation status throughout folliculogenesis was observed, we hypothesized that reactive oxygen species-induced cellular damage could potentially already start to accumulate from the earliest stages of folliculogenesis in dictyate arrested oocytes residing in primordial follicles.

### Increased ROS-induced protein and lipid damage in primordial follicles of advanced maternal age

Since reactive oxygen species can already be created during the primordial follicle stage, we further investigated whether ROS-induced damage was already detectable at the primordial follicle stage and thus could contribute to ovarian ageing. For this, we visualized ROS-induced damage at the protein, lipid and DNA level (Lim and Luderer, 2011). To identify a possible age-dependent trend in ROS-induced damage, a linear regression analysis was carried out for the maximum intensity of the different antibodies and female age. Protein oxidation was visualized using antibodies against 3-nitrotyrosine (NTY) (Figure 2A), and significantly increased in oocytes with increasing female age (Figure 2B). The same is true for lipid peroxidation, which was visualized using antibodies against 4-hydroxynonenal (4HNE) (Figure 2C and D). In contrast, DNA oxidization, visualized using an antibody against 8-Oxo-2’-deoxyguanosine (8OHDG) (Figure 2E), did not differ between different female ages (Figure 2F). Negative controls showed no staining for ROS induced antibodies (Figure S1B-D). Together, these data indicated the accumulation of oxidative damage to proteins and lipids in oocytes of advanced maternal age already at the primordial follicle stage.

**Figure 2.**
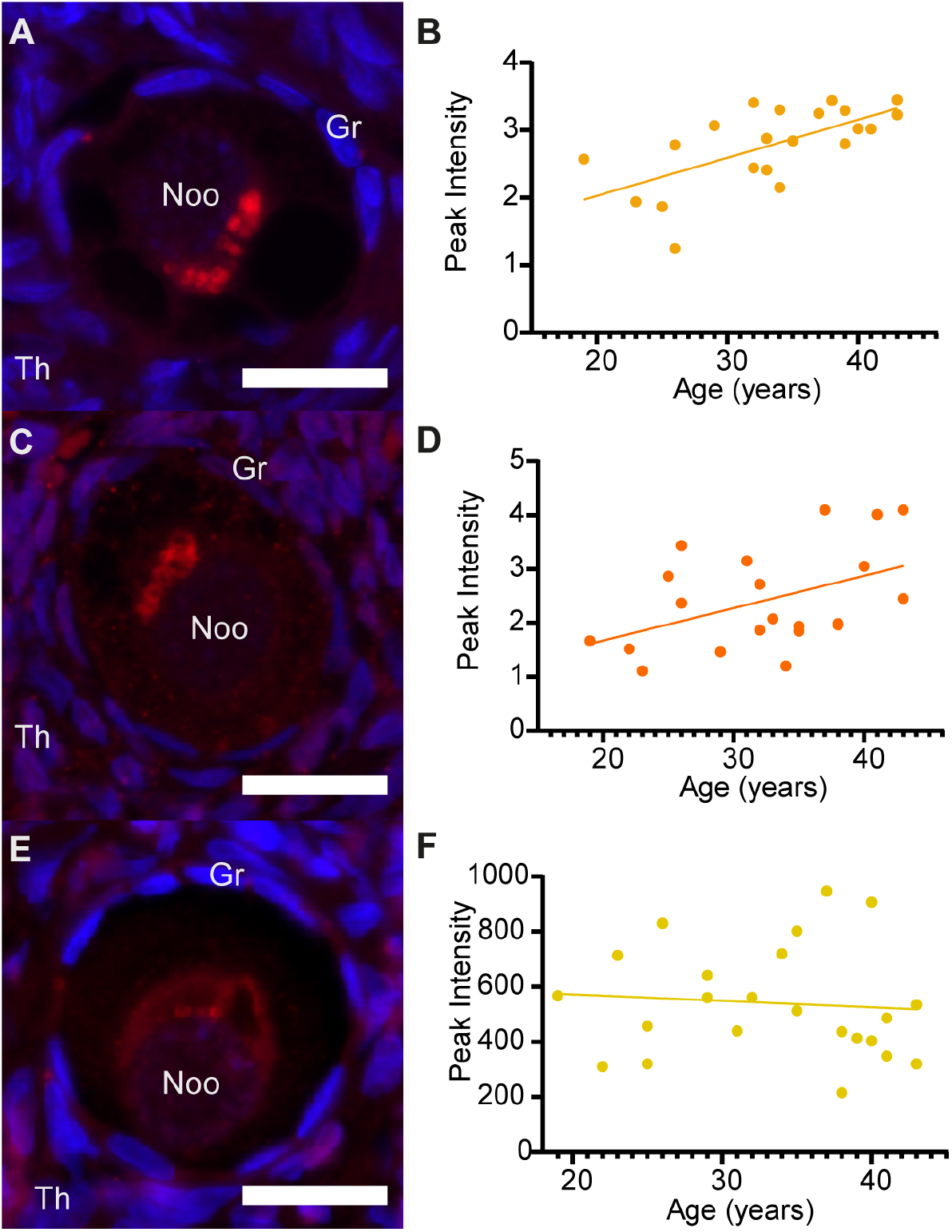
Reactive oxygen species (ROS) induced damage to proteins and lipids increases with age in primordial follicles. Immunofluorescence staining of different ROS damage markers in primordial follicles in human ovarian tissue and the simple linear regressions of the peak intensity of these markers with age in human oocytes. (A and B) ROS-induced protein oxidation-staining (mouse anti 3-nitrotyrosine) significantly increased with increasing female age, R^2^ = 0.4092, p = 0.002 (C and D) ROS-induced lipid peroxidation-staining (mouse anti 4-hydroxynonenal) significantly increased with increasing female age, R^2^ = 0.2052, p = 0.0392. (E and F) ROS-induced DNA damage-staining (anti 8-Oxo-2’-deoxyguanosine) did not change with increasing female age, R^2^ = 0.0073, p = 0.6983. Scale bars = 20μm. Noo = nucleus oocyte, Gr = granulosa cells, Th = theca cells. Blue = DAPI, Red = ROS induced damage.

### Targeted metabolomics and lipidomics suggest impaired mitochondrial function in oocytes of advanced maternal age

Since we found higher ROS-induced damage to lipids and proteins in aged primordial follicles, we questioned whether cellular metabolism could be impaired in aged oocytes. Therefore, we performed metabolomics and lipidomics by UPLC-mass spectrometry on GV oocytes, MI oocytes and cumulus cells associated with these oocytes. The oocytes were collected after ovarian hyper stimulation for Intra Cytoplasmic Sperm Injection (ICSI) treatment or oocyte preservation treatment, but appeared to be immature so could not be used for treatment. To achieve sufficient yield for subsequent metabolomics and lipidomics, ten oocytes were grouped by age per analysis. Because one ICSI treatment yielded sufficient amounts of cumulus cells, cumulus cell samples did not need to be pooled. Age categories were chosen based on the oocyte quality decline observed with ovarian ageing (Schieve et al., 2003) and characterized as follows: (1) donor age: 23-34 years old, representing good quality oocytes; (2) Donor age: 35-37 years old, oocytes with a gradual decline in quality, (3) Donor age: 38-42 years old, reflecting rapid decline in oocyte quality. MI oocytes were categorized in two age groups, as follows: (1) 23-34 years old, and (2) 35-42 years old. All oocytes were studied in triplicate per category, thus thirty oocytes were included per category, 150 in total.

Using metabolomics and lipidomics, we detected and annotated 89 polar metabolites and over 1100 lipid species. The metabolome (Figure 3A and B) and lipidome (Figure S2A and B) of human GV and MI oocytes showed a distinct pattern with increasing female age as identified by partial least squares discriminant analysis (PLS-DA) (Figure 3A and B). From this PLS-DA analysis, metabolites with a *variable of importance in projection* (VIP) score >1 are considered significant and are shown in Figure 3D. We performed the same statistical analysis to compare the youngest to the oldest age groups of GV oocytes (Figure S3). For lipidome analysis, we quantified changes of lipid classes; i.e., the sum of lipids belonging to one class, rather than individual lipids, using one way ANOVA, student’s t-test or a Kruskal Wallis test (p-value of < 0.05 was considered significant).

**Figure 3.**
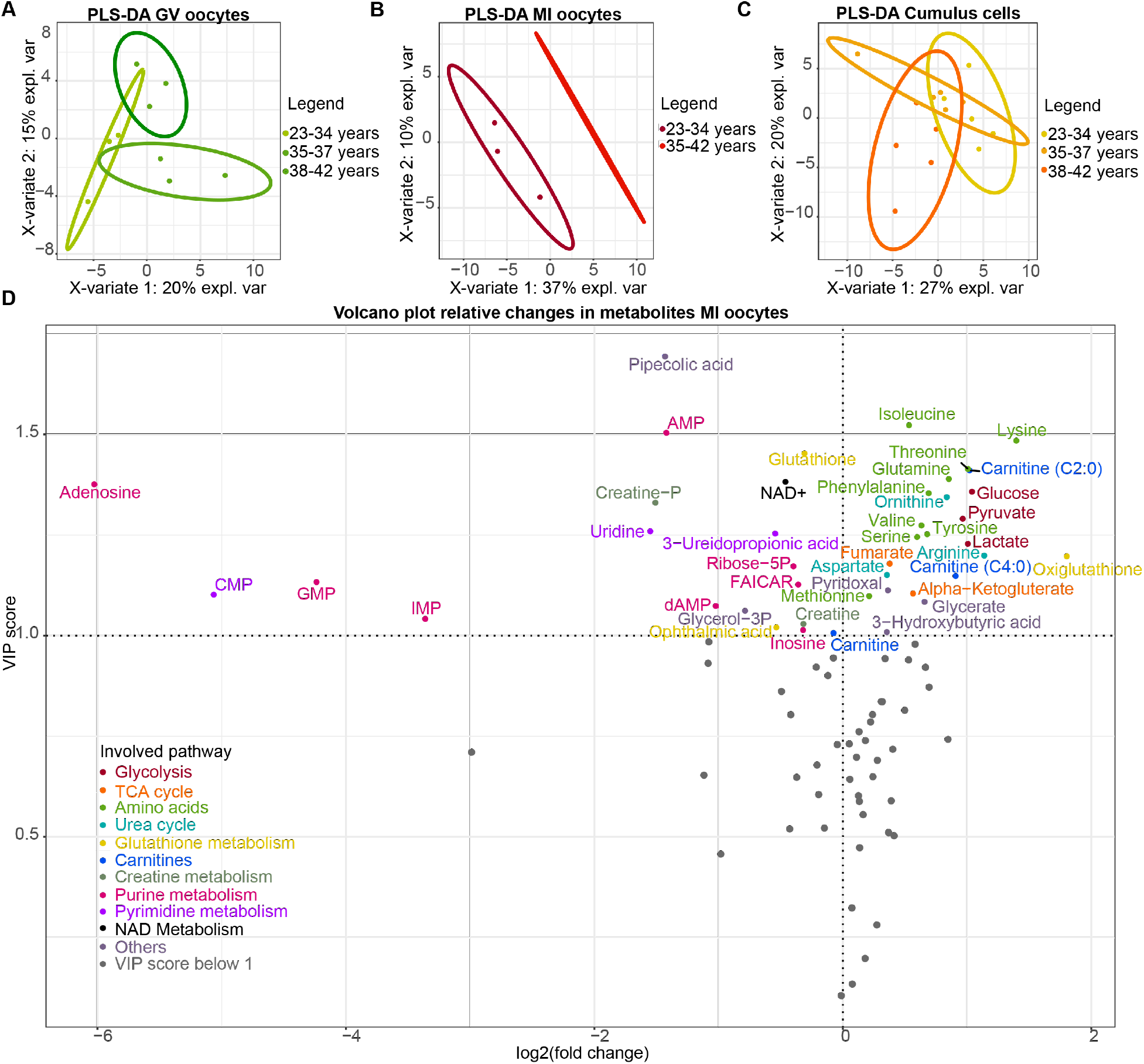
Age-related metabolite changes in MI and GV oocytes. (A) PLS-DA analysis of the metabolome of pooled human GV oocytes distinguishes between different female age categories: 23-34 years, 35-37 years and 38-42 years. (B) PLS-DA analysis of the metabolome of pooled human MI oocytes distinguishes between different female age categories: 23-34 years and 35-42 years. (C) PLS-DA analysis of the metabolome of single cumulus cell samples per female age category: 23-34 years, 35-37 years and 38-42 years. There was no clear distinction between different age categories. (D) volcano plot of relative changes in metabolites of MI oocytes between different age categories based on VIP scores > 1, colored by pathway in which they play the most important role.

In cumulus cells, PLS-DA analysis of both the metabolome and lipidome could not distinguish different age groups (Figure 3C, Figure S2C).

### Increased signs of oxidative stress in ageing GV and MI oocytes

Since we found an age-related increase in ROS-induced damage in primordial follicles, we investigated several indicators for oxidative stress through metabolomics in GV and MI oocytes. One of these indicators is the glutathione to oxiglutathione ratio (GSH/GSSG) within a cell. In GV oocytes, glutathione did not change with age and oxigluthathione decreased (Figure 4A and B. In contrast to GV, a decrease in glutathione abundance was observed for older MI oocytes, which was accompanied with an apparent increase of oxiglutathione (Figure 4A and B). This indicates a shift towards GSSG and thus increased oxidative stress in MI oocytes of advanced maternal age.

**Figure 4.**
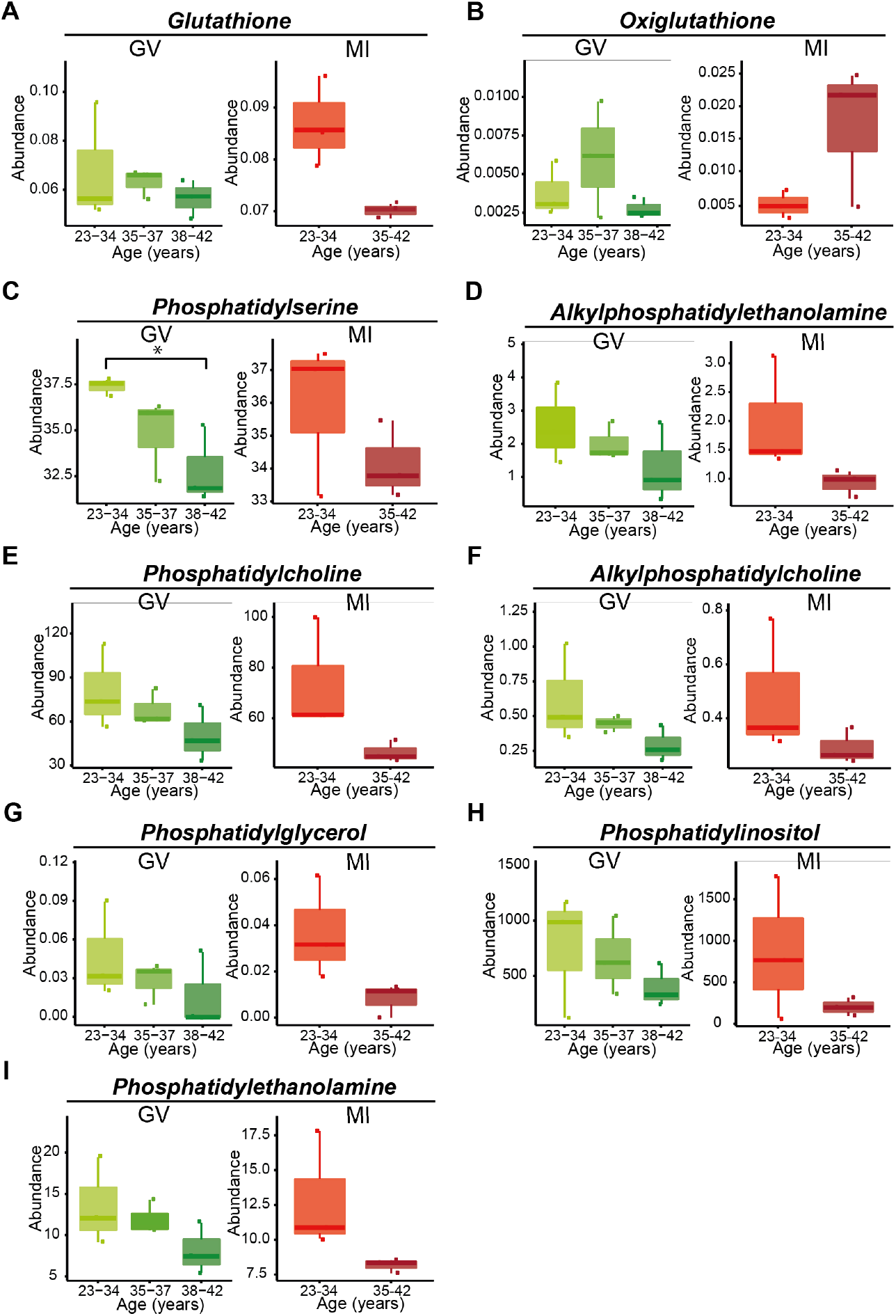
Different signs of oxidative stress are seen in GV and MI oocytes. (A and B) Glutathione is depleted in aging oocytes during MI, while oxiglutathione accumulates, indicating an increase in oxidative stress. (C and D) Phosphatidylserine significantly decreased with age in GV oocytes and showed a decreasing trend in MI oocytes. Alkylphosphatidylethanolamine showed a decreasing trend with age in GV and MI oocytes (E-I) Phosphatidylcholine, Alkylphosphatidylcholine, Phosphatidylglycerol and Phosphatidylinositol, and Phosphatidylethanolamine show a decreasing trend with age, both in GV and MI oocytes. GV = GV oocytes, MI = MI oocytes.

Another indicator for oxidative damage is lipid peroxidation. Lipid peroxidation was already visualized in primordial follicles of women of older ovarian age (Figure 2B). Using lipidomics, we investigated age-related changes in different lipid classes in GV and MI oocytes. To our surprise, all phospholipids decreased in abundance with increasing age in both GV and MI oocytes (Figure 4C-I). In GV, a significant age-related decrease was observed in total phosphatidylserine (PS) and a decreasing trend was seen for phosphatidylcholine (PC), phosphatidylethanolamine (PE), phosphatidylglycerols (PG) and to lesser extent phosphatidylinositols (PI) (Figure 4C-I). In MI, we found a decreasing trend for PE, PC, PG and PI (Figure 4C-I). In addition to lipid classes, many individual lipids changed with increasing female age, Figure S4 shows an overview of different individual lipid changes with female age in MI oocytes and all data on individual lipid changes can be found in the Table S1D.

### Metabolites feeding into the TCA cycle accumulate with increasing age in both GV and MI oocytes

The TCA cycle, located in the mitochondria, is mostly known for, but is not limited to the most important supplier of adenosine triphosphate (ATP) (Akram, 2014). In both GV and MI oocytes, metabolites associated with glycolysis increased. In GV oocytes, glucose and pyruvate increased with increasing female age (Figure 5A and B) In MI oocytes, glucose, glucose-6-phosphate, phosphoenolpyruvate, pyruvate and lactate, showed an increase in oocytes of advanced maternal age, of which glucose and pyruvate roughly doubled (Figure 5A-C). In contrast, metabolites involved in glycolysis did not change with age in cumulus cell samples (Table S1C).

**Figure 5.**
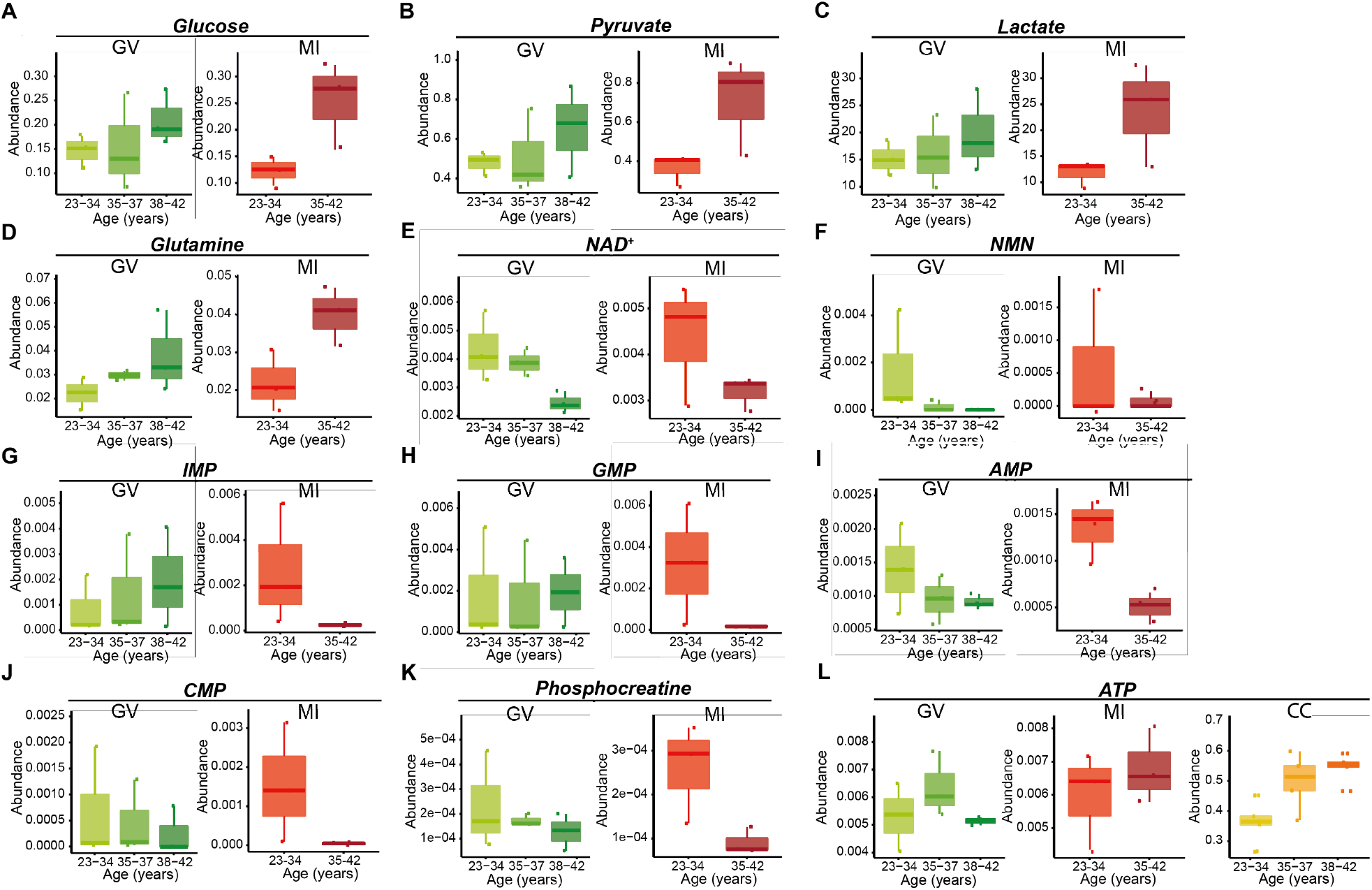
Metabolites associated with mitochondrial function are disturbed in aging oocytes, especially in MI oocytes. (A-D) Metabolites that fuel mitochondrial metabolism accumulate with age: glucose, pyruvate, lactate and glutamine. (E and F) Metabolites associated with energy status of the cell diminish in aging oocyte: NAD^+^ and NMN. (G-J) Monophosphate nucleotide intermediates, that are dependent on mitochondrial metabolism, show depletion with age: IMP, GMP, AMP, CMP (K) Phosphocreatine together with AMP is associated with the adenosine salvage pathway, an alternative path to creating ATP, both reduce with age. (L) ATP increases in cumulus cells with age; possibly in an attempt to compensate for mitochondrial dysfunction in the aging oocyte. GV = GV oocytes, MI = MI oocytes, CC = Cumulus cells.

In addition to pyruvate, glutamine functions as mitochondrial fuel. Glutamine also accumulated with age in both GV and MI oocytes (Figure 5D). TCA cycle intermediates, such as α-ketoglutarate, succinate and fumarate did not change with age in MI oocytes (Table S1B), while the TCA cycle intermediates succinate and fumarate were decreased in GV oocytes (Figure S3). The fact that substrates feeding into the TCA cycle accumulate with age, while TCA cycle intermediates did not change in MI oocytes, might be indicative of the TCA cycle not being able to process available substrates.

### Impaired nicotinamide adenine dinucleotide (NAD^+^) biosynthesis in older GV and MI oocytes

NAD^+^ plays an important role in energy metabolism (Houtkooper et al., 2010, Zapata-Pérez et al., 2021, Katsyuba et al., 2020) and stimulates mitochondrial biogenesis through SIRT1 (Brenmoehl and Hoeflich, 2013). In GV oocytes, we observed a decrease in NAD^+^ and nicotinamide mononucleotide (NMN, an NAD^+^ precursor) abundance (Figure 5E-F), while other NAD^+^ precursors, kynurenine and nicotinamide riboside (NR), increased with age (Figure S3). The downstream product of kynurenine, quinolinic acid, did not accumulate in GV oocytes (Table S1A). In MI oocytes, NAD^+^ also decreased in abundance with age, but the precursors NMN, kynurenine and NR all remained unchanged throughout the different age categories (Figure 5E and F, Table S1B). Interestingly, nicotinamide riboside (NR) was higher with age in cumulus cells (Table S1C). An age-related decrease in NAD^+^ in both GV and MI oocytes could indicate that NAD^+^ biosynthesis is impaired in oocytes of women of advanced maternal age. This is specifically true in GV oocytes, since several NAD^+^ precursors accumulated with age, but do not result in an increase in NAD^+^.

### Changes in mitochondrial purine and pyrimidine nucleotide biosynthesis in older MI oocytes

Purine and pyrimidine nucleotide biosynthesis, is largely carried out by mitochondria and provides the cell with nucleotides (Wang, 2016). The purine nucleotide-synthesis pathway requires a significant amount of molecular fuel. For each molecule of inosine monophosphate (IMP) that is generated, five molecules of ATP are required, as well as glutamine, glycine, aspartate, formate and carbon dioxide (Chan et al., 2018). Many of these substrates are produced directly by the mitochondria and it has recently been shown that a large proportion of the purinosomes are grouped together and co-localize with mitochondria (Chan et al., 2018, Yin et al., 2018). In GV oocytes, glycine, and its precursor serine, increased with age (Table S1A). In MI oocytes, glycine abundance did not change with age, but the precursor serine showed a 1.5-fold increase (Table S1B). As mentioned before, glutamine concentration accumulated with age in both GV and MI oocytes (Figure 5D). Aspartate accumulated in both GV and MI oocytes (Table S1A and B). The direct precursor to the de novo purine synthesis pathway, ribose-5P, is produced in the cytosol and did not show an age-dependent trend in GV and was decreased in MI oocytes of advanced maternal age. Dramatic differences were observed in the monophosphate end-products at the core of this pathway; an almost 10-fold decrease in inosine monophosphate (IMP), a 20-fold decrease in guanosine monophosphate (GMP) and a 2.7-fold decrease in adenosine monophosphate (AMP) were observed in MI oocytes of advanced maternal age (Figure 4G, H, I). In GV oocytes, AMP also decreased with increasing age, but IMP increased and GMP did not change with female age (Table S1A). Decreases were also seen in MI oocytes for 5-formamidoimidazole-4-carboxamide ribotide (FAICAR), inosine and adenosine (Table S1B). This was not the case for GV oocytes, in which adenosine also decreased, but inosine increased and FAICAR did not change in women of advanced maternal age (Table S1A). In cumulus cells metabolites of the purine nucleotide biosynthesis were not affected with age (Table S1C).

In line with observations in purine metabolism, monophosphate end-products of pyrimidine synthesis were also affected strongly in older MI oocytes. Older MI oocytes showed a 33-fold lower abundance in cytidine monophosphate (CMP), a 1.5-fold decrease for the uracil breakdown product 3-ureidopropionic acid, and a 3-fold lower abundance of uridine (Figure 5J, Table S1B). In GV oocytes and cumulus cells, no age-dependent changes were observed in pyrimidine synthesis (Table S1A and S1C). Together, these results show that the mitochondrial contribution to purine and pyrimidine nucleotide biosynthesis appears highly disrupted with age in MI oocytes.

### Alternative energy sources in aged GV and MI oocytes

As mentioned before, glycolysis intermediates seemed to be increased in GV and MI oocytes of women of advanced maternal age (Figure 5A - C). Besides increasing glycolysis, oocytes are able to use the adenosine salvage pathway as another alternative energy source to the TCA cycle, for which they use phosphocreatine and AMP (Scantland et al., 2014). AMP was decreased in GV oocytes and in MI oocytes of advanced maternal age (Figure 5I). GV oocytes and cumulus cells did not show a decrease in phosphocreatine, but in MI oocytes phosphocreatine showed a 2.7-fold decrease (Figure 5K, Table S1B). Creatine was also decreased with age in MI oocytes (not in GV oocytes), but only 1.2-fold (Table S1B).

It has been previously described that oocytes are supplied with ATP by surrounding cumulus cells through gap junctions (Dalton et al., 2014, Van Blerkom et al., 2008). Therefore, cumulus cells might partly compensate for the loss of ATP due to mitochondrial dysfunction (Kansaku et al., 2017). We observed an increase in ATP levels of cumulus cells with age, while the ATP levels of GV and MI oocytes seemed unchanged (Figure 5L).

## Discussion

It is often assumed that mitochondria in dictyate arrested oocytes in primordial follicles are largely inactive to prevent ROS-induced oxidative stress. However, increasing evidence indicates that mitochondria in dictyate arrested oocytes are already metabolically active (Bradley and Swann, 2019, Van Blerkom, 2011). Recently, Wang et al., 2020 studied the transcriptomic landscape of ovarian cell types of cynomolgus monkeys during different stages of folliculogenesis and found that already during early stages of folliculogenesis antioxidant proteins were downregulated, while oxidative stress markers were upregulated in oocytes of older monkeys (Wang et al., 2020). Using high resolution fluorescence microscopy, we studied the activation status of the mitochondrial gatekeeper enzyme PDH on human ovarian biopsies of women of different ages. We did not find significant differences in PDH activation status during different stages of folliculogenesis or with age, indicating that mitochondria are indeed active in dictyate arrested oocytes in primordial follicles. In addition, by using immunofluorescence markers, we have shown an age-related increase in lipid peroxidation and protein oxidation, clear signs of oxidative damage, in dictyate arrested oocytes at the primordial follicle stage. We did not observe an age-related increase in DNA oxidation. This could mean that ageing oocytes are somehow protected against DNA oxidation. Alternatively, and perhaps more likely, oocytes with excessive DNA damage may have disappeared from the ovarian reserve and thus not observed in this study. Signs of oxidative damage were also observed in metabolomic and lipidomic analysis of human GV and MI oocytes, leftover oocytes from ICSI and oocyte preservation treatments: in aged MI oocytes we found a disturbed glutathione to oxiglutathione ratio and in both MI and GV oocytes, we found a major drop in phospholipids, as will be discussed further below. Besides signs of oxidative stress, we also observed age-related changes that indicate mitochondrial dysfunction: impaired NAD^+^, purine and pyrimidine pathways and the use of alternative energy sources, such as the adenosine salvage pathway and increased ATP in surrounding cumulus cells.

We found several metabolic changes that all pointed towards mitochondrial dysfunction in MI and, to lesser extent, GV oocytes of advanced maternal age. GV oocytes are transcriptionally active, but during their development to metaphase II oocytes, they become transcriptionally silent, which means that they cannot replace degrading gene transcripts anymore (Ntostis et al., 2021, Bouniol-Baly et al., 1999). Therefore, age-induced damage is expected to have a more pronounced effect on MI oocytes compared to GV oocytes (Llonch et al., 2021). An increase in glycolytic metabolites in women of older age could point towards an affected TCA cycle but could also point towards an increase in anaerobic glycolysis. Since both pyruvate and glutamine are increased as well, it is more likely that the TCA cycle is unable to process the provided metabolites. The observed impaired NAD^+^ biosynthesis may for example be due to a lack of nucleotides in older oocytes or may point towards a broader dysfunction in the mitochondria (Imai and Guarente, 2014). Besides, a decrease in NAD^+^ can lead to decreased mitochondrial biogenesis, inducing a downward spiral (Katsyuba et al., 2020). MI oocytes of advanced maternal age also showed large decreases in mitochondria-derived substrates and end products, e.g., inosine monophosphate (IMP), guanosine monophosphate (GMP), and cytidine monophosphate (CMP), of the purine and pyrimidine nucleotide biosynthesis. Since we observed an age-related increase in metabolites related to glycolysis, a decrease in the metabolites of the adenosine salvage pathway, and an increase in ATP content of surrounding cumulus cells, we hypothesize that these sources could be used as alternative energy sources in oocytes of advanced maternal age. Besides this, the changes with age in GSH/GSSG ratio in MI and GV oocytes, point towards an increased state of oxidative stress in oocytes of advanced maternal age (Zitka et al., 2012). Glutathione itself is involved in numerous oocyte and embryo functions. For example, by protecting the meiotic spindle against oxidative stress, but also by stimulating sperm decondensation required for the development of the male pronucleus after fertilization (Luberda, 2005, Zuelke et al., 1997, Yoshida et al., 1993, Sutovsky and Schatten, 1997). Depletion of the glutathione pool is therefore expected to have adverse effects on the oocyte. Moreover, almost all phospholipid classes in older oocytes in both the GV and MI stage showed a decreasing trend with age. It should be noted that this effect does not necessarily mean an absolute reduction in the total abundance of each class. It seems plausible that products of these affected phospholipids, for example oxidized phospholipids, are not detected using the current method; either because they fall below detection limits or are turned into products that are not annotated. Using an immunofluorescence antibody against 4-hydroxynonenal (another form of lipid peroxidation), we have already found that oocytes of ovarian biopsies of women of advanced maternal age suffer more lipid peroxidation. Oxidation of the phospholipids of advanced maternal age oocytes also seems in line with the extremely disturbed GSH:GSSG ratio. Phospholipids contain a relatively large number of poly unsaturated fatty acids (PUFAs). PUFAs are more sensitive to this type of destruction than saturated classes, as radicals specifically attack double bonds (Hu et al., 2017, Catalá, 2009). In line with this, de la Barca et al., 2017, found decreased polyunsaturated choline plasmalogens in follicular fluid of women with diminished ovarian reserve, which they also explain by an increase in oxidative stress (de la Barca et al., 2017). Previous studies into aging and oxidative stress in mouse oocytes showed the vulnerability of phospholipids to lipid peroxidation, already at the GV stage of development (Mok et al., 2016). Besides having a function as signaling molecules, phospholipids are found mostly on the cell membrane (Catalá, 2009, Shimada and Terada, 2001), where they are responsible for the structure, and thereby permeability, of the membrane. Therefore, the lower amount of phospholipids may by itself already have a detrimental effect on oocyte function, for example by decreasing the chance of successful fertilization in women of advanced maternal age.

In the current study, we used comprehensive metabolomic and lipidomic techniques on human GV, MI oocytes and cumulus cells. While some research has been published on the use of metabolomics and lipidomics in reproductive medicine, this concerned non-invasive measurements that were done on culture medium or follicular fluid (Siristatidis et al., 2018, Montani et al., 2019, de la Barca et al., 2017, Bouet et al., 2020, Luti et al., 2020). Although interesting from a clinical perspective, these studies could not unravel the biological mechanisms that play a role in oocyte quality or ovarian ageing, since these studies did not measure metabolites in oocytes or cumulus cells. Additionally, previous lipidomic studies were limited in scope, focusing on a small selection of lipids using e.g. MALDI or MRM (Zhang et al., 2020, Cataldi et al., 2013), rather than perform full-scan LC-MS analysis of complex lipids as we have done here, which provides a much more comprehensive picture of the oocyte and cumulus cell lipidomes.

The current study provides evidence that mitochondrial dysfunction is one of the important mechanisms underlying ovarian ageing. Mechanisms that could improve mitochondrial function in oocytes are therefore interesting to investigate in future studies. Mitochondrial replacement therapy by ooplasm transfer could be a potential treatment for advanced maternal age. However, ooplasm transfer includes medical risks and ethical objections and is therefore not yet a good treatment option for advanced maternal age (Yamada et al., 2021). Other possible treatment options include enhancing the depleted NAD^+^ pool or correcting glutathione deficiency in oocytes of advanced maternal age. NAD^+^ promotes mitochondrial biogenesis (Brenmoehl and Hoeflich, 2013). In mice, NMN, an NAD precursor that can boost NAD^+^, has already been shown to improve oocyte quality (Bertoldo et al., 2020), and recently a pilot clinical trial was published in which glycine and N-acetylcysteine showed to improve glutathione deficiency and mitochondrial function (Kumar et al., 2021). Therefore, supplementing embryo medium of advanced maternal age embryos with these compounds could be a treatment option worth exploring.

To achieve sufficient yield for the metabolomics and lipidomics of oocytes, ten oocytes were pooled per analysis. Due to the limited amounts of oocytes available, only three replicates per age group were possible. Therefore, most results did not reach statistical significance, based on a p value of < 0.05 and after correction for multiple testing. Still, the total amount of oocytes included in this study is 150, which is much larger than the number of oocytes used for most human studies so far (Trebichalská et al., 2021, Llonch et al., 2021, Yuan et al., 2021) and, combined with 39 human ovarian biopsies, makes this a unique dataset.

Ideally, we would have additionally included metaphase II oocytes in our study, but these oocytes are all intended for use in ICSI treatments. GV and MI oocytes are also collected during oocyte pick up but, due to immaturity, cannot be used for ICSI treatment, and could therefore be used in our study. It is important to note that results might have been different when metaphase II oocytes, and especially when naturally ovulated oocytes, would have been included. The fact that oocytes used in the current study were all subjected to ovarian hyperstimulation with high doses of follicle stimulating hormones, might have concealed some age-related differences.

## Methods

### Ethical approval

The protocol of the study was discussed by the local Medical Ethical Committee and it was decided that the study adhered to national guidelines.

### Experimental model

Ovarian tissue was collected through PALGA: “the nationwide network and registry of histo- and cytopathology in the Netherlands” (Casparie, et al., 2007). We collected ovarian tissue stored in the Netherlands after surgery between 1991 and 2018. Ovarian tissue samples were fixed in 10% formalin, embedded in paraffin and divided into different age groups: 18-29, 30-34, 35-40 and 41-45 years. A tissue sample was excluded in case the pathology report included: malignancies, polycystic ovarian syndrome or preventive resection due to BRCA mutation carrier. Further patient characteristics were not provided. A total of 39 ovarian tissue samples of individual patients were assessed.

150 immature oocytes (90 GV oocytes and 60 MI oocytes) (not suitable for ICSI as they were immature) and 15 cumulus cell samples of women undergoing ICSI treatment based on male or tubal factor infertility or fertility preservation for non-medical reasons in the Amsterdam UMC, location Academic Medical Center, were included in this study. Women were excluded if they, or their partners, were HIV or hepatitis B/C positive. For metabolomics and lipidomics on human oocytes, 10 oocytes per analysis were pooled. Each analysis was repeated three times. Women were divided in three age groups: 23-34, 35-37 and 38-42 years of age for GV oocytes; and two age groups for MI oocytes: 23-34 and 35-42 years. Cumulus cell samples were not pooled and each analysis was repeated five times.

### Follicle identification using Haematoxylin Eosin (HE)

To determine whether follicles were present in the received tissues, tissues were cut into 5-μm-thick sections. For dewaxing and hydration, sections were treated with xylene (VWR chemicals), after which they were exposed to an ethanol gradient. The sections were rinsed in distilled water for three minutes and stained with hematoxylin (Roth) for five minutes. After staining, the sections were rinsed twice shortly with distilled water for half a minute and placed under running tap water for ten minutes. Subsequently, the sections were stained with eosin (VWR chemicals) in 70% ethanol for at least three minutes. The sections were then shortly rinsed, dehydrated using an ethanol-increasing gradient and xylene and mounted using Entellan (Sigma-Aldrich).

### Immunofluorescence of mitochondria and oxidative phosphorylation use

When at least one follicular stage was identified in an ovarian section, immunofluorescence staining was performed. Again, tissues were cut into 5-μm-thick sections, dewaxed and hydrated, after which they were shortly washed in distilled water. For antigen retrieval, sections were heated in 0,01M sodium citrate until boiling for 10 minutes in a microwave and afterwards placed under running tap water for 10 minutes. All slides were subsequently washed in TBS and incubated with 100 μl Superblock (Scytek Laboratories) at room temperature for one hour, to prevent a-specific binding of antibodies. The slides were incubated overnight at 4 degrees Celsius with primary antibodies or 200μg/ml Mouse IgG (Vector Labs) and 200μg/ml Rabbit IgG (Vector Labs) for negative controls. Primary antibodies used in this study were 5μg/ml Mouse monoclonal anti-Pyruvate Dehydrogenase E1-alpha subunit antibody (Abcam) and 10μg/ml Rabbit polyclonal phosphoDetect™ PDHE1α pSer^293^ (PDHA) (Merck). On the second day, the slides were washed three times for 10 minutes with TBS including 0,1% Tween-20 and incubated with secondary antibodies: 2 mg/ml Abberior STAR fluorescence 635p, Goat anti Mouse (Abberior) and 2 mg/ml Abberior STAR fluorescence 580, Goat anti Rabbit (Abberior) for 1 hour. Subsequently, the slides were washed with TBS and incubated with 1 mg/ml 4’,6-diamidino-2-phenylindole (DAPI) (Sigma) for 5 minutes. For cell surface visualization, the fluorescent lectin wheat germ agglutinin (WGA) (Thermo Fisher) was used in a concentration of 1 mg/ml in TBS for 5 minutes. Lastly, the slides were washed with TBS, embedded with Prolong © Gold Antifade Reagent and placed at 4 degrees Celsius.

Z-stack images of follicles at different developmental stages were taken on a Leica SP8 fluorescence microscope with stimulated emission depletion (STED). Depending on follicle size, a 100x or 40x oil immersion lens with a Z-stack step size of 0,15 μm with a total of 7 steps using Leica Application Suite X. Settings remained unchanged during follicle imaging. All images underwent deconvolution with Huygens Professional version 18.04.

**T**o measure oxidative damage, immunofluorescence was carried out as described above using 2μg/ml Mouse anti-8-Hydroxy-2’-deoxyguanosine (JaICA) visualizing DNA oxidation, 4μg/ml Mouse anti-4-Hydroxynonenal Monoclonal Antibody (Thermo Fisher) visualizing lipid peroxidation or 10 μg/ml Mouse Anti-3-Nitrotyrosine antibody (Abcam) visualizing nitrated proteins. For negative controls, Mouse IgG (Vector Labs) in the same concentration as the antibodies were used. As secondary antibody, 2μg/ml fluorescence AF555 (Invitrogen) was used. DAPI and WGA were used as counterstaining. All sections were stained at the same time in order to minimize technical variety. Images were taken on a Leica D5000B Microscope with 40x air objective.

Image analysis was done on images after deconvolution using Leica Application Suite X. Mean immunofluorescence intensity of the different channels representing different antibody stainings was measured by drawing a region of interest (ROI) around the oocyte and surrounding cumulus cells. Immunofluorescence intensity ratios were determined for the PDHE and PDHE1α pSer^293^ antibodies by dividing the immunofluorescence intensity of PDHE by the immunofluorescence intensity of PDHE1α pSer^293^ (PDHE: PDHE1α pSer^293^) and were calculated and compared between the different stages of folliculogenesis and between the different age groups. IBM SPSS Statistics 25.0 was used for statistical analysis. Differences between developmental stages and age groups were calculated using the Mann-Whitney U-test. Differences were considered significant in case p<0.05.

Fluorescence intensity of the different oxidative damage antibodies was measured using Leica Application Suite AF. Differences in fluorescence intensity between age groups were calculated using the Mann-Whitney U-test with IBM SPSS Statistics 25.0. Differences were considered significant in case of p-value <0.05.

### Targeted metabolomics and lipidomics on human oocytes and cumulus cells

Metabolomics and lipidomics were performed as previously described, with minor adjustments (Schomakers et al., 2022). For each sample, approximately 150.000 cumulus cells or exactly 10 oocytes were added to a 2 mL Eppendorf Safe-Lock tube. In the same 2 mL tube, the following internal standards dissolved in water were added for metabolomics:adenosine-^15^N_5_-monophosphate (5 nmol), adenosine-^15^N_5_-triphosphate (5 nmol), D_4_-alanine (0.5 nmol), D_7_-arginine (0.5 nmol), D_3_-aspartic acid (0.5 nmol), D_3_-carnitine (0.5 nmol), D_4_-citric acid (0.5 nmol), ^13^C_1_-citrulline (0.5 nmol), ^13^C_6_-fructose-1,6-diphosphate (1 nmol), ^13^C_2_-glycine (5 nmol), guanosine-^15^N_5_-monophosphate (5 nmol), guanosine-^15^N_5_-triphosphate (5 nmol), ^13^C_6_-glucose (10 nmol), ^13^C_6_-glucose-6-phosphate (1 nmol), D_3_-glutamic acid (0.5 nmol), D_5_-glutamine (0.5 nmol), D_5_-glutathione (1 nmol), ^13^C_6_-isoleucine (0.5 nmol), D_3_-lactic acid (1 nmol), D_3_-leucine (0.5 nmol), D_4_-lysine (0.5 nmol), D_3_-methionine (0.5 nmol), D_6_-ornithine (0.5 nmol), D_5_-phenylalanine (0.5 nmol), D_7_-proline (0.5 nmol), ^13^C_3_-pyruvate (0.5 nmol), D_3_-serine (0.5 nmol), D_6_-succinic acid (0.5 nmol), D_5_-tryptophan (0.5 nmol), D_4_-tyrosine (0.5 nmol), D_8_-valine (0.5 nmol). In the same 2 mL tube, the following amounts of internal standards dissolved in 1:1 (v/v) methanol:chloroform were added for Lipidomics: Bis(monoacylglycero)phosphate BMP(14:0)2 (0.2 nmol), Ceramide-1-phosphate C1P (d18:1/12:0) (0.125 nmol), D_7_-Cholesteryl Ester CE(16:0) (2.5 nmol), Ceramide Cer(d18:1/12:0) (0.125 nmol), Ceramide Cer(d18:1/25:0) (0.125 nmol), Cardiolipin CL(14:0)4 (0.1 nmol), Diacylglycerol DAG(14:0)2 (0.5 nmol), Glucose Ceramide GlcCer(d18:1/12:0) (0.125 nmol), Lactose Ceramide LacCer(d18:1/12:0) (0.125 nmol), Lysophosphatidicacid LPA(14:0) (0.1 nmol), Lysophosphatidylcholine LPC(14:0) (0.5 nmol), Lysophosphatidylethanolamine LPE(14:0) (0.1 nmol), Lysophosphatidylglycerol LPG(14:0) (0.02 nmol), Phosphatidic acid PA(14:0)2 (0.5 nmol), Phosphatidylcholine PC(14:0)2 (2 nmol), Phosphatidylethanolamine PE(14:0)2 (0.5 nmol), Phosphatidylglycerol PG(14:0)2 (0.1 nmol), Phosphatidylinositol PI(8:0)2 (0.5 nmol), Phosphatidylserine PS(14:0)2 (5 nmol), Sphinganine 1-phosphate S1P(d17:0) (0.125 nmol), Sphinganine-1-phosphate S1P(d17:1) (0.125 nmol), Ceramide phosphocholines SM(d18:1/12:0) (2.125 nmol), Sphingosine SPH(d17:0) (0.125 nmol), Sphingosine SPH(d17:1) (0.125 nmol), Triacylglycerol TAG(14:0)2 (0.5 nmol). After adding internal standards, solvents were added to each sample, for a total of 500 μL water, 500 μL methanol and 1 mL of chloroform. Samples were then thoroughly mixed, followed by centrifugation for 10 minutes at 14,000 rpm, creating a two-phase system with protein precipitate in the middle.

#### Metabolomics

The top layer, containing the polar phase, was transferred to a new 1.5 mL tube and dried using a vacuum concentrator at 60°C. Dried samples were reconstituted in 50 μL 6:4 (v/v) methanol:water. Metabolites were analyzed using a Waters Acquity ultra-high performance liquid chromatography system coupled to a Bruker Impact II™ Ultra-High Resolution Qq-Time-Of-Flight mass spectrometer. Samples were kept at 12°C during analysis and 5 μL of each sample was injected. Injection order was randomized, with a pooled QC sample measured at the start, end, and between every ten samples. Chromatographic separation was achieved using a Merck Millipore SeQuant ZIC-cHILIC column (PEEK 100 × 2.1 mm, 3 μm particle size). Column temperature was held at 30°C. Mobile phase consisted of (A) 1:9 (v/v) acetonitrile:water and (B) 9:1 (v/v) acetonitrile:water, both containing 5 mmol/L ammonium acetate. Using a flow rate of 0.25 mL/min, the LC gradient consisted of: 100% B for 0-2 min, reach 0% B at 28 min, 0% B for 28-30 min, reach 100% B at 31 min, 100% B for 31-32 min. Column re-equilibration is achieved by increasing the flow rate to 0.4 mL/min at 100% B for 32-35 min. MS data were acquired using both positive and negative ionization in full scan mode over the range of m/z 50-1200. Data were analyzed using Bruker TASQ software version 2.1.22.3. All reported metabolite intensities were normalized to the number of oocytes for oocyte data, or the total adenosine nucleotide (TAN) pool for cumulus cells, as well as to internal standards with comparable retention times and response in the MS. Metabolite identification has been based on a combination of accurate mass, (relative) retention times and fragmentation spectra, compared to the analysis of a library of standards.

#### Lipidomics

The bottom layer of the extraction, containing the apolar phase, was transferred to a glass vial and evaporated under a stream of nitrogen at 60°C. The residue was dissolved in 100 μL of 1:1 (v/v) methanol:chloroform. Lipids were analyzed using a Thermo Scientific Ultimate 3000 binary HPLC coupled to a Q Exactive Plus Orbitrap mass spectrometer. For normal phase separation, 2 μL of each sample was injected onto a Phenomenex^®^ LUNA silica, 250 * 2 mm, 5μm 100Å. Injection order was randomized, with a pooled QC sample measured at the start, end, and between every ten samples. Column temperature was held at 25°C. Mobile phase consisted of (A) 85:15 (v/v) methanol:water containing 0.0125% formic acid and 3.35 mmol/L ammonia and (B) 97:3 (v/v) chloroform:methanol containing 0.0125% formic acid. Using a flow rate of 0.3 mL/min, the LC gradient consisted of: 10% A for 0-1 min, reach 20% A at 4 min, reach 85% A at 12 min, reach 100% A at 12.1 min, 100% A for 12.1-14 min, reach 10% A at 14.1 min, 10% A for 14.1-15 min. For reversed phase separation, 5 μL of each sample was injected onto a Waters HSS T3 column (150 x 2.1 mm, 1.8 μm particle size). Column temperature was held at 60°C. Mobile phase consisted of (A) 4:6 (v/v) methanol:water and B 1:9 (v/v) methanol:isopropanol, both containing 0.1% formic acid and 10 mmol/L ammonia. Using a flow rate of 0.4 mL/min, the LC gradient consisted of: 100% A at 0 min, reach 80% A at 1 min, reach 0% A at 16 min, 0% A for 16-20 min, reach 100% A at 20.1 min, 100% A for 20.1-21 min. MS data were acquired using negative and positive ionization using continuous scanning over the range of m/z 150 to m/z 2000. Data were analyzed using a lipidomics pipeline developed in-house, written in the R programming language (http://ww.r-project.org). All reported lipids were normalized to corresponding internal standards according to lipid class, as well as to the number of oocytes for oocyte data, or the total adenosine nucleotide (TAN) pool for cumulus cells. Lipid identification has been based on a combination of accurate mass, (relative) retention times and the injection of relevant standards.

All data generated with metabolomics and lipidomics can be found in the Supplementary data. For the metabolomics, a partial least square regression discriminant analysis (PLS-DA) was used to discriminate between the different age groups using the programming language R and the mixOmics package. Differential changes of metabolites and lipids contributing to the discrimination between the different age groups, were determined according to the Variable Importance in Projection (VIP) score extracted from the PLS-DA. Since different age groups showed a distinct pattern in the PLS-DA analysis of different age groups for the MI and GV oocytes, metabolites with a VIP score of >1 were considered significant. Since the PLS-DA analysis of cumulus cells could not distinguish different age groups, differences were calculated using one-way ANOVA, with a significance cut off of < 0.05. PLS-DA analysis was also able to clearly distinct age groups in MI and GV oocytes. Age-related changes for individual lipids can be found in the supplementary data. To investigate age-related changes in lipid classes rather than individual lipids, p-values were calculated by a student’s t-test, one-way ANOVA or Kruskal Wallis test, depending on normality of distribution. P-value of <0.05 was considered significant.

## Supporting information

Supplementary data_Smits et al 2023

## Acknowledgements

The authors acknowledge the Center for Reproductive Medicine Fertility Laboratory for providing the GV and MI oocytes and cumulus cells for research; PALGA: Dutch Pathology Registry for using human ovarian biopsies (Casparie et al., 2007); Milou Kleijn and Stella Dorrestein for their help optimizing the immunofluorescence assays; Cindy Korver and Saskia van Daalen for running the immunofluorescence assays for ROS-induced damage; Ronald Wanders for critically reading the manuscript; the Amsterdam Reproduction & Development research institute for financial support (grant number V.000296).

## Authors’ Contributions

M.A.J.S., B.V.S., M. van W., R.C.I.W., S.M., M.G., R.H.H. and G.H. designed the project, M.A.J.S. collected ovarian tissue and oocytes and cumulus cell samples, M.A.J.S. performed immunofluorescence experiments, B.V.S. and E.J.M.W. performed metabolomic and lipidomic experiments, M.A.J.S., B.V.S., E.J.M.W and G.E.J. analyzed data and drafted figures. M.A.J.S., B.V.S. and G.H. wrote the manuscript. All authors have reviewed the manuscript.

## Data availability statement

The published article includes all metabolomics and lipidomics datasets on GV, MI and cumulus cell samples generated or analyzed during this study.

---

**Figure S1.**
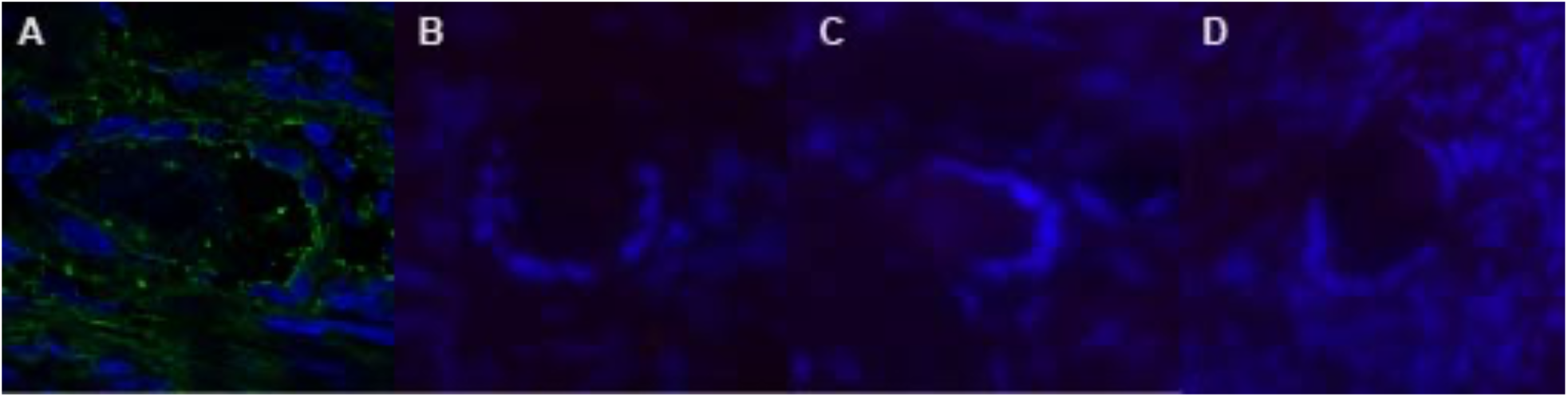
IgGs of immunofluorescence staining ovarian biopsies. Negative controls of ovarian biopsies used in this manuscript. Instead of antibody, mouse and/or rabbit IgG was used in the same concentration as the primary antibody. (A) Negative control of figure 1E-H. Primary antibodies used were Mouse IgG (red) and Rabbit IgG (yellow). Slides were also incubated with DAPI (blue), and wheat germ agglutinin (green). (B) Negative control for figure 2A. Red = mouse IgG, blue = DAPI (C) Negative control for figure 2B. Red = mouse IgG, blue = DAPI (D) Negative control for figure 2C. Red = mouse IgG, blue = DAPI

**Figure S2.**
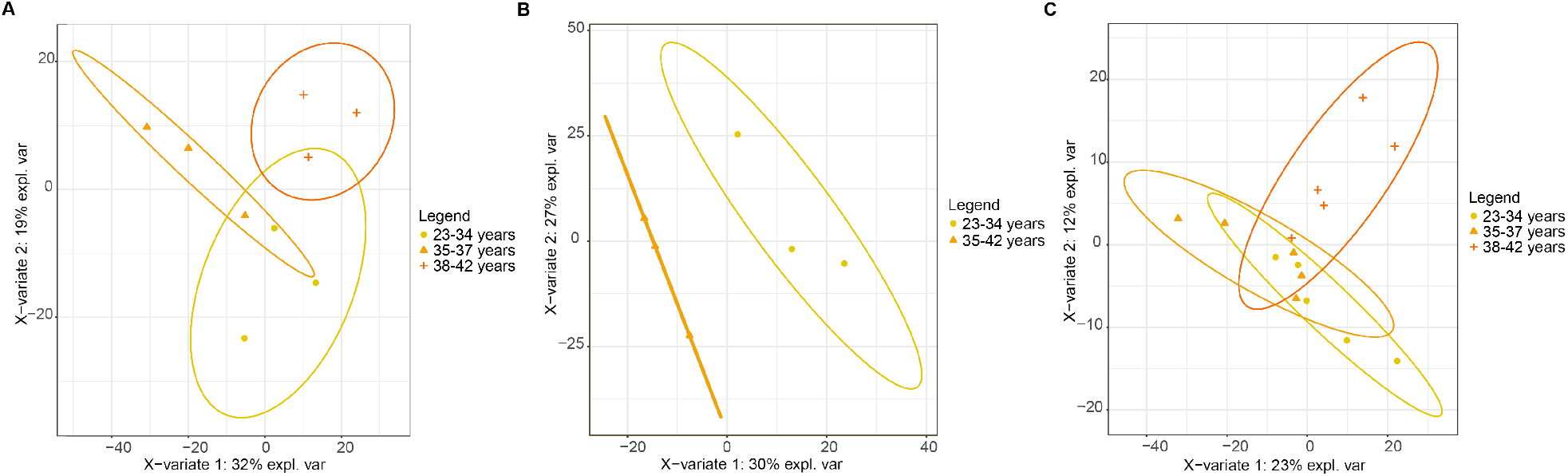
2 PLS-DA analysis distinguishes different age groups in GV and MI oocytes, but not in cumulus cells. (A) PLS-DA analysis of the lipidome of pooled human GV oocytes distinguishes between female age categories: 23-34 years, 35-37 years and 38-42 years. (B) PLS-DA analysis of the lipidome of pooled human MI oocytes distinguishes between female age categories: 23-34 years and 35-42 years. (C) PLS-DA analysis of the lipidome of single cumulus cell samples per female age category: 23-34 years, 35-37 years and 38-42 years. There was no clear distinction between different age categories.

**Figure S3.**
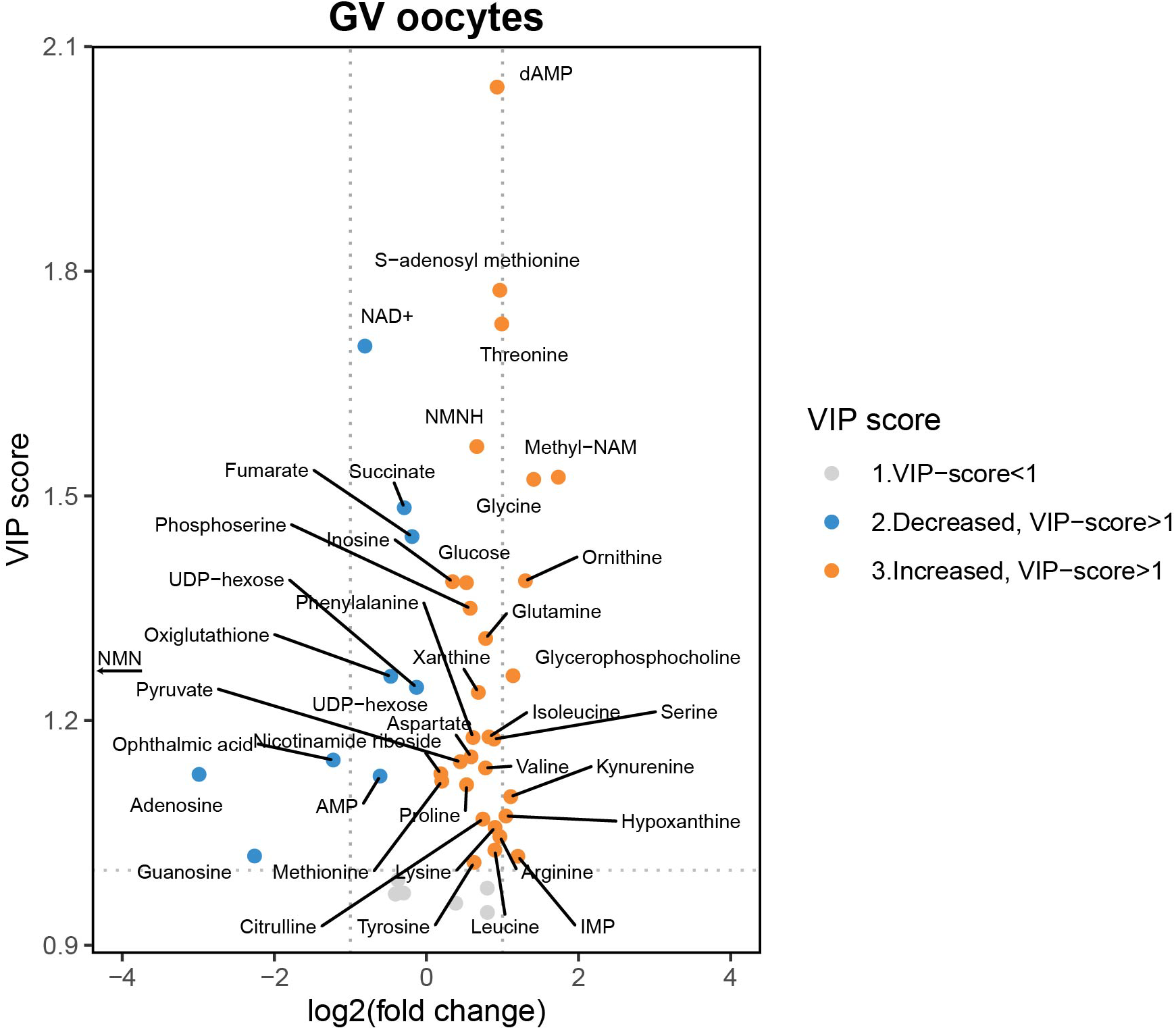
Volcano plot of age-related changes between GV oocytes of young (23-34) and advanced maternal age (38-42) based on a VIP score > 1. Volcano plot of relative changes in metabolites of GV oocytes between the youngest (23-34 years) and oldest (38-42 years) age group, based on VIP scores > 1. Note: NMN levels were below detection limits in the oldest age group, making it impossible to calculate an accurate log2(fold change) and VIP score. Nevertheless, it is included in the graph, as an age related decrease below detection limits is highly relevant.

**Figure S4.**
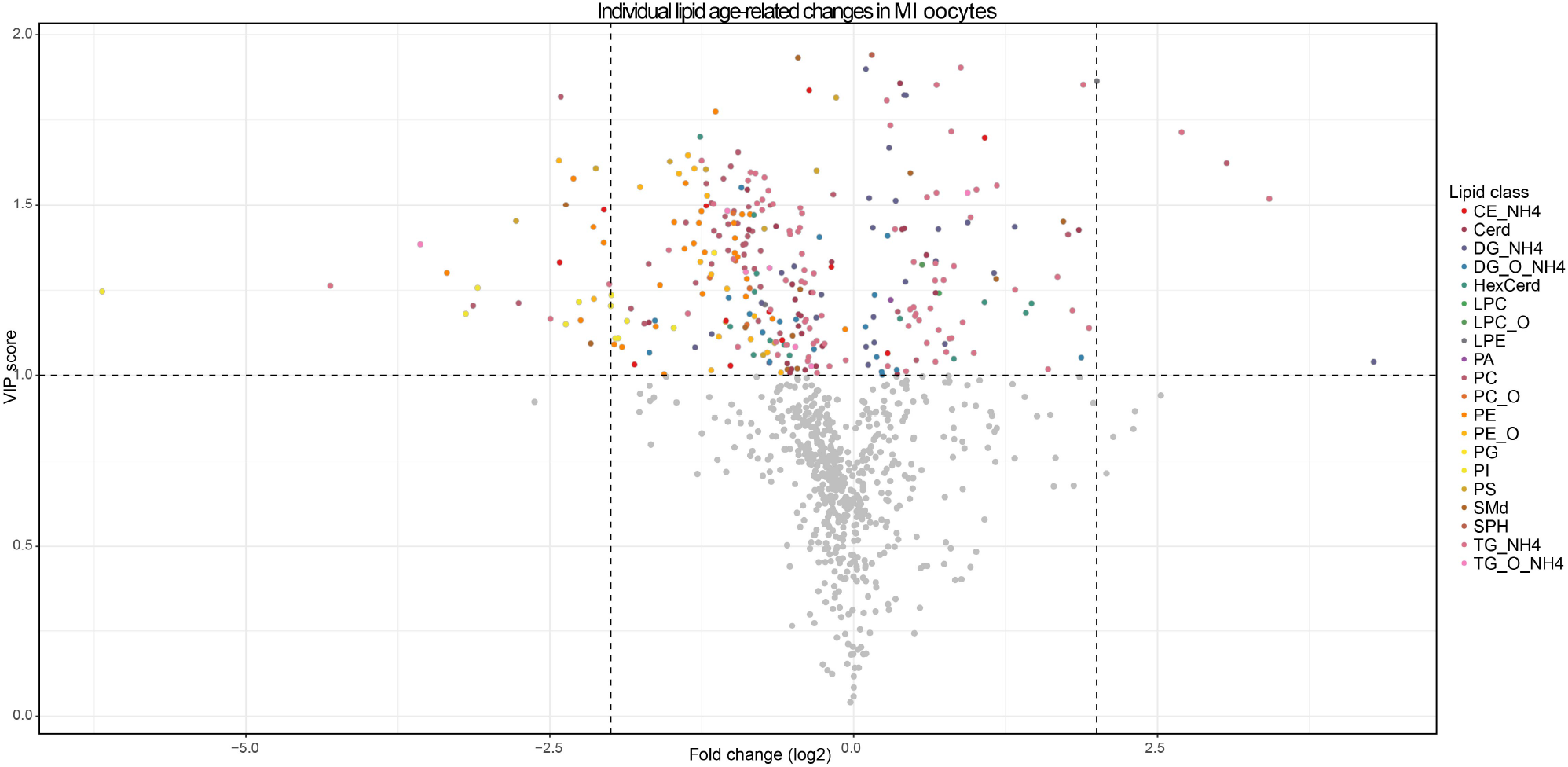
many age-related lipids changed with increasing female age in MI oocytes. Volcano plot of relative changes in individual lipids of MI oocytes between the youngest (23-34 years) and oldest (35-42 years) age group, based on VIP scores > 1. Individual lipids are coloured based on the lipid class they belong to.

## References

Ahmed, T. A., Ahmed, S. M., El-Gammal, Z., Shouman, S., Ahmed, A., Mansour, R., and El-Badri, N. 2019. Oocyte Aging: The Role of Cellular and Environmental Factors and Impact on Female Fertility. Adv Exp Med Biol.

Akram, M. 2014. Citric acid cycle and role of its intermediates in metabolism. Cell Biochem Biophys, 68, 475–8.

Bertoldo, M. J., Listijono, D. R., Ho, W. J., Riepsamen, A. H., Goss, D. M., Richani, D., Jin, X. L., Mahbub, S., Campbell, J. M., Habibalahi, A., et al. 2020. NAD(+) Repletion Rescues Female Fertility during Reproductive Aging. Cell Rep, 30, 1670–1681.e7.

Bouet, P. E., Boueilh, T., de la Barca, J. M. C., Boucret, L., Blanchard, S., Ferré-L’Hotellier, V., Jeannin, P., Descamps, P., Procaccio, V., Reynier, P., et al. 2020. The cytokine profile of follicular fluid changes during ovarian ageing. J Gynecol Obstet Hum Reprod, 49, 101704.

Bouniol-Baly, C., Hamraoui, L., Guibert, J., Beaujean, N., Szöllösi, M. S., and Debey, P. 1999. Differential transcriptional activity associated with chromatin configuration in fully grown mouse germinal vesicle oocytes. Biol Reprod, 60, 580–7.

Bradley, J., and Swann, K. 2019. Mitochondria and lipid metabolism in mammalian oocytes and early embryos. Int J Dev Biol, 63, 93–103.

Brenmoehl, J., and Hoeflich, A. 2013. Dual control of mitochondrial biogenesis by sirtuin 1 and sirtuin 3. Mitochondrion, 13, 755–61.

Casparie, M., Tiebosch, A. T., Burger, G., Blauwgeers, H., van de Pol, A., van Krieken, J. H., and Meijer, G. A. 2007. Pathology databanking and biobanking in The Netherlands, a central role for PALGA, the nationwide histopathology and cytopathology data network and archive. Cell Oncol, 29, 19–24.

Catalá, A. 2009. Lipid peroxidation of membrane phospholipids generates hydroxy-alkenals and oxidized phospholipids active in physiological and/or pathological conditions. Chem Phys Lipids, 157, 1–11.

Cataldi, T., Cordeiro, F. B., Costa Ldo, V., Pilau, E. J., Ferreira, C. R., Gozzo, F. C., Eberlin, M. N., Bertolla, R. P., Cedenho, A. P., and Turco, E. G. 2013. Lipid profiling of follicular fluid from women undergoing IVF: young poor ovarian responders versus normal responders. Hum Fertil (Camb), 16, 269–77.

CDC 2016. 2016 National Summary ART sucess rates. Center for Disease Control and prevention.

Chan, C. Y., Pedley, A. M., Kim, D., Xia, C., Zhuang, X., and Benkovic, S. J. 2018. Microtubule-directed transport of purine metabolons drives their cytosolic transit to mitochondria. Proceedings of the National Academy of Sciences, 115, 13009–13014.

Chiang, J. L., Shukla, P., Pagidas, K., Ahmed, N. S., Karri, S., Gunn, D. D., Hurd, W. W., and Singh, K. K. 2020. Mitochondria in Ovarian Aging and Reproductive Longevity. Ageing Res Rev, 63, 101168.

Cinco, R., Digman, M. A., Gratton, E., and Luderer, U. 2016. Spatial Characterization of Bioenergetics and Metabolism of Primordial to Preovulatory Follicles in Whole Ex Vivo Murine Ovary. Biol Reprod, 95, 129.

Dalton, C. M., Szabadkai, G., and Carroll, J. 2014. Measurement of ATP in single oocytes: impact of maturation and cumulus cells on levels and consumption. J Cell Physiol, 229, 353–61.

de la Barca, J. M. C., Boueilh, T., Simard, G., Boucret, L., Ferré-L’Hotellier, V., Tessier, L., Gadras, C., Bouet, P. E., Descamps, P., Procaccio, V., et al. 2017. Targeted metabolomics reveals reduced levels of polyunsaturated choline plasmalogens and a smaller dimethylarginine/arginine ratio in the follicular fluid of patients with a diminished ovarian reserve. Hum Reprod, 32, 2269–2278.

Eijkemans, M. J., van Poppel, F., Habbema, D. F., Smith, K. R., Leridon, H., and te Velde, E. R. 2014. Too old to have children? Lessons from natural fertility populations. Hum Reprod, 29, 1304–12.

Faddy, M. J., Gosden, R. G., Gougeon, A., Richardson, S. J., and Nelson, J. F. 1992. Accelerated disappearance of ovarian follicles in mid-life: implications for forecasting menopause. Hum Reprod, 7, 1342–6.

Hashimoto, S., Morimoto, N., Yamanaka, M., Matsumoto, H., Yamochi, T., Goto, H., Inoue, M., Nakaoka, Y., Shibahara, H., and Morimoto, Y. 2017. Quantitative and qualitative changes of mitochondria in human preimplantation embryos. J Assist Reprod Genet, 34, 573–580.

Hekimi, S., Lapointe, J., and Wen, Y. 2011. Taking a “good” look at free radicals in the aging process. Trends Cell Biol, 21, 569–76.

Houtkooper, R. H., Cantó, C., Wanders, R. J., and Auwerx, J. 2010. The secret life of NAD+: an old metabolite controlling new metabolic signaling pathways. Endocr Rev, 31, 194–223.

Hu, C., Wang, M., and Han, X. 2017. Shotgun lipidomics in substantiating lipid peroxidation in redox biology: Methods and applications. Redox biology, 12, 946–955.

Imai, S., and Guarente, L. 2014. NAD+ and sirtuins in aging and disease. Trends Cell Biol, 24, 464–71.

Jansen, R. P., and de Boer, K. 1998. The bottleneck: mitochondrial imperatives in oogenesis and ovarian follicular fate. Mol Cell Endocrinol, 145, 81–8.

Kansaku, K., Itami, N., Kawahara-Miki, R., Shirasuna, K., Kuwayama, T., and Iwata, H. 2017. Differential effects of mitochondrial inhibitors on porcine granulosa cells and oocytes. Theriogenology, 103, 98–103.

Kasapoğlu, I., and Seli, E. 2020. Mitochondrial Dysfunction and Ovarian Aging. Endocrinology, 161.

Katsyuba, E., Romani, M., Hofer, D., and Auwerx, J. 2020. NAD+ homeostasis in health and disease. Nature Metabolism, 2, 9–31.

Kirkwood, T. B. 1998. Ovarian ageing and the general biology of senescence. Maturitas, 30, 105–11.

Kumar, P., Liu, C., Hsu, J. W., Chacko, S., Minard, C., Jahoor, F., and Sekhar, R. V. 2021. Glycine and N-acetylcysteine (GlyNAC) supplementation in older adults improves glutathione deficiency, oxidative stress, mitochondrial dysfunction, inflammation, insulin resistance, endothelial dysfunction, genotoxicity, muscle strength, and cognition: Results of a pilot clinical trial. Clin Transl Med, 11, e372.

Lefkimmiatis, K., Grisan, F., Iannucci, L. F., Surdo, N. C., Pozzan, T., and Di Benedetto, G. 2020. Mitochondrial communication in the context of aging. Aging Clin Exp Res.

Lim, J., and Luderer, U. 2011. Oxidative damage increases and antioxidant gene expression decreases with aging in the mouse ovary. Biol Reprod, 84, 775–82.

Llonch, S., Barragán, M., Nieto, P., Mallol, A., Elosua-Bayes, M., Lorden, P., Ruiz, S., Zambelli, F., Heyn, H., Vassena, R., et al. 2021. Single human oocyte transcriptome analysis reveals distinct maturation stage-dependent pathways impacted by age. Aging Cell, 20, e13360.

Luberda, Z. 2005. The role of glutathione in mammalian gametes. Reprod Biol, 5, 5–17.

Luti, S., Fiaschi, T., Magherini, F., Modesti, P. A., Piomboni, P., Governini, L., Luddi, A., Amoresano, A., Illiano, A., Pinto, G., et al. 2020. Relationship between the metabolic and lipid profile in follicular fluid of women undergoing in vitro fertilization. Mol Reprod Dev, 87, 986–997.

May-Panloup, P., Boucret, L., Chao de la Barca, J. M., Desquiret-Dumas, V., Ferre-L’Hotellier, V., Moriniere, C., Descamps, P., Procaccio, V., and Reynier, P. 2016. Ovarian ageing: the role of mitochondria in oocytes and follicles. Hum Reprod Update, 22, 725–743.

Mok, H. J., Shin, H., Lee, J. W., Lee, G. K., Suh, C. S., Kim, K. P., and Lim, H. J. 2016. Age-Associated Lipidome Changes in Metaphase II Mouse Oocytes. PLoS One, 11, e0148577.

Montani, D. A., Braga, D., Borges, E., Jr., Camargo, M., Cordeiro, F. B., Pilau, E. J., Gozzo, F. C., Fraietta, R., and Lo Turco, E. G. 2019. Understanding mechanisms of oocyte development by follicular fluid lipidomics. J Assist Reprod Genet, 36, 1003–1011.

Ntostis, P., Iles, D., Kokkali, G., Vaxevanoglou, T., Kanavakis, E., Pantou, A., Huntriss, J., Pantos, K., and Picton, H. M. 2021. The impact of maternal age on gene expression during the GV to MII transition in euploid human oocytes. Human Reproduction.

Pasquariello, R., Ermisch, A. F., Silva, E., McCormick, S., Logsdon, D., Barfield, J. P., Schoolcraft, W. B., and Krisher, R. L. 2019. Alterations in oocyte mitochondrial number and function are related to spindle defects and occur with maternal aging in mice and humansdagger. Biol Reprod, 100, 971–981.

Polonio, A. M., Chico-Sordo, L., Córdova-Oriz, I., Medrano, M., García-Velasco, J. A., and Varela, E. 2020. Impact of Ovarian Aging in Reproduction: From Telomeres and Mice Models to Ovarian Rejuvenation. Yale J Biol Med, 93, 561–569.

Scantland, S., Tessaro, I., Macabelli, C. H., Macaulay, A. D., Cagnone, G., Fournier, É., Luciano, A. M., and Robert, C. 2014. The adenosine salvage pathway as an alternative to mitochondrial production of ATP in maturing mammalian oocytes. Biol Reprod, 91, 75.

Schieve, L. A., Tatham, L., Peterson, H. B., Toner, J., and Jeng, G. 2003. Spontaneous abortion among pregnancies conceived using assisted reproductive technology in the United States. Obstet Gynecol, 101, 959–67.

Schomakers, B. V., Hermans, J., Jaspers, Y. R. J., Salomons, G., Vaz, F. M., van Weeghel, M., and Houtkooper, R. H. 2022. Polar metabolomics in human muscle biopsies using a liquid-liquid extraction and full-scan LC-MS. STAR Protoc, 3, 101302.

Shimada, M., and Terada, T. 2001. Phosphatidylinositol 3-kinase in cumulus cells and oocytes is responsible for activation of oocyte mitogen-activated protein kinase during meiotic progression beyond the meiosis I stage in pigs. Biol Reprod, 64, 1106–14.

Siristatidis, C. S., Sertedaki, E., Vaidakis, D., Varounis, C., and Trivella, M. 2018. Metabolomics for improving pregnancy outcomes in women undergoing assisted reproductive technologies. Cochrane Database Syst Rev, 3, Cd011872.

Sutovsky, P., and Schatten, G. 1997. Depletion of glutathione during bovine oocyte maturation reversibly blocks the decondensation of the male pronucleus and pronuclear apposition during fertilization. Biol Reprod, 56, 1503–12.

Trebichalská, Z., Kyjovská, D., Kloudová, S., Otevřel, P., Hampl, A., and Holubcová, Z. 2021. Cytoplasmic maturation in human oocytes: an ultrastructural study †. Biol Reprod, 104, 106–116.

Van Blerkom, J. 2011. Mitochondrial function in the human oocyte and embryo and their role in developmental competence. Mitochondrion, 11, 797–813.

Van Blerkom, J., Davis, P., and Thalhammer, V. 2008. Regulation of mitochondrial polarity in mouse and human oocytes: the influence of cumulus derived nitric oxide. Molecular Human Reproduction, 14, 431–444.

Vasileiou, P. V. S., Evangelou, K., Vlasis, K., Fildisis, G., Panayiotidis, M. I., Chronopoulos, E., Passias, P. G., Kouloukoussa, M., Gorgoulis, V. G., and Havaki, S. 2019. Mitochondrial Homeostasis and Cellular Senescence. Cells, 8.

Wang, C. H., Wu, S. B., Wu, Y. T., and Wei, Y. H. 2013. Oxidative stress response elicited by mitochondrial dysfunction: implication in the pathophysiology of aging. Exp Biol Med (Maywood), 238, 450–60.

Wang, L. 2016. Mitochondrial purine and pyrimidine metabolism and beyond. Nucleosides Nucleotides Nucleic Acids, 35, 578–594.

Wang, L., Tang, J., Wang, L., Tan, F., Song, H., Zhou, J., and Li, F. 2021. Oxidative stress in oocyte aging and female reproduction. J Cell Physiol, 236, 7966–7983.

Wang, S., Zheng, Y., Li, J., Yu, Y., Zhang, W., Song, M., Liu, Z., Min, Z., Hu, H., Jing, Y., et al. 2020. Single-Cell Transcriptomic Atlas of Primate Ovarian Aging. Cell, 180, 585–600.e19.

Wang, T., Zhang, M., Jiang, Z., and Seli, E. 2017. Mitochondrial dysfunction and ovarian aging. Am J Reprod Immunol, 77.

Wilding, M., Dale, B., Marino, M., di Matteo, L., Alviggi, C., Pisaturo, M. L., Lombardi, L., and De Placido, G. 2001. Mitochondrial aggregation patterns and activity in human oocytes and preimplantation embryos. Human Reproduction, 16, 909–917.

Wilding, M., De Placido, G., De Matteo, L., Marino, M., Alviggi, C., and Dale, B. 2003. Chaotic mosaicism in human preimplantation embryos is correlated with a low mitochondrial membrane potential. Fertil Steril, 79, 340–6.

Yamada, M., Sato, S., Ooka, R., Akashi, K., Nakamura, A., Miyado, K., Akutsu, H., and Tanaka, M. 2021. Mitochondrial replacement by genome transfer in human oocytes: Efficacy, concerns, and legality. Reprod Med Biol, 20, 53–61.

Yin, J., Ren, W., Huang, X., Deng, J., Li, T., and Yin, Y. 2018. Potential Mechanisms Connecting Purine Metabolism and Cancer Therapy. Frontiers in Immunology, 9.

Yoshida, M., Ishigaki, K., Nagai, T., Chikyu, M., and Pursel, V. G. 1993. Glutathione concentration during maturation and after fertilization in pig oocytes: relevance to the ability of oocytes to form male pronucleus. Biol Reprod, 49, 89–94.

Yuan, L., Yin, P., Yan, H., Zhong, X., Ren, C., Li, K., Heng, B. C., Zhang, W., and Tong, G. 2021. Single-cell transcriptome analysis of human oocyte ageing. J Cell Mol Med, 25, 6289–303.

Zapata-Pérez, R., Wanders, R. J. A., van Karnebeek, C. D. M., and Houtkooper, R. H. 2021. NAD(+) homeostasis in human health and disease. EMBO Mol Med, 13, e13943.

Zhang, X., Wang, T., Song, J., Deng, J., and Sun, Z. 2020. Study on follicular fluid metabolomics components at different ages based on lipid metabolism. Reprod Biol Endocrinol, 18, 42.

Zitka, O., Skalickova, S., Gumulec, J., Masarik, M., Adam, V., Hubalek, J., Trnkova, L., Kruseova, J., Eckschlager, T., and Kizek, R. 2012. Redox status expressed as GSH:GSSG ratio as a marker for oxidative stress in paediatric tumour patients. Oncology letters, 4, 1247–1253.

Zuelke, K. A., Jones, D. P., and Perreault, S. D. 1997. Glutathione oxidation is associated with altered microtubule function and disrupted fertilization in mature hamster oocytes. Biol Reprod, 57, 1413–9.

